# Mechanical wounding impacts the growth versus defence balance in tomato (*Solanum lycopersicum*)

**DOI:** 10.1101/2022.11.24.517841

**Authors:** Ana Flavia Aparecida Cunha, Pedro Henrique Duarte Rodrigues, Ana Clara Anghinoni, Vinicius Juliani de Paiva, Daniel Gonçalves da Silva Pinheiro, Marcelo Lattarulo Campos

## Abstract

Plants have evolved elaborate surveillance systems that allow them to perceive the attack by pests and pathogens and activate the appropriate defences. Mechanical stimulation, such as mechanical wounding, represents one of the most reliable cues for the perception of potential herbivore aggressors. Here we demonstrate that mechanical wounding disturbs the growth versus defence balance in tomato, a physiological condition where growth reduction arises as a pleiotropic consequence of the activation of defence responses (or vice-versa). We observed that consecutive lesions on tomato leaves impairs the formation of several growth-related traits, including shoot elongation, leaf expansion and time for flowering set, while concomitantly activating the production of defence responses such as trichome formation and the upregulation of defence-related genes. We also provide genetic evidence that this wound-induced growth repression is a consequence of tomato plants sensing the injuries via jasmonates (JAs), a class of plant hormones known to be master regulators of the plant growth versus defence balance. Besides providing a mechanistic explanation on how the growth and defence balance is shifted when plants are subjected to a specific type of mechanical stimulus, our results may offer a practical explanation for why tomato productivity is so negatively impacted by herbivore attack.

**Highlight:** Antagonism between growth and defence responses was observed in tomato plants subject to mechanical wounding, a treatment that hinders development while promoting the activation of anti-herbivore traits.

## Introduction

Plants are constantly challenged by a myriad of pest and pathogens that utilize the green tissues as a source for nutritional needs. To survive, plants have evolved elaborate surveillance systems that allow them to perceive the potential aggressor and trigger the appropriate defence responses. In this sense, mechanical stimulation represents one of the most consistent cues for recognition of attacking herbivores (Waterman et al., 2019; Kollasch et al., 2020; Matsumara et al., 2022). When subjected to mechanical stimulation, such as that caused by arthropod movement or tissue injury, plants can activate diverse signalling cascades that culminate in the activation of multiple defence barriers, including the induction of trichomes, the synthesis of toxic metabolites and the production of proteinase inhibitors (PIs) (Green and Ryan, 1972; Traw and Bergelson, 2003; Howe and Jander, 2008; Mostafa et al., 2022). It is also recognized that mechanosensory stimuli can trigger large modifications in plant transcriptional activity, leading to the upregulation of numerous defence-related genes (Heidel-Fischer, 2014; Matsumara et al., 2022). However, due to the diversity and complexity of this kind of environmental signal, several gaps remain in our knowledge to understand the molecular framework utilized by plants to sense specific types of mechanical stimulation and what are the overall consequences of its perception to plant development besides the activation of defence responses.

Mechanical wounding is a simple and reproductible technique to study the consequences of mechanical stimulation in plants. This procedure, which is commonly referred as “simulated herbivory”, consists of using razors or other mechanical means to produce similar damage patterns in the plant tissues as those caused by arthropod herbivores (Baldwin, 1990; Zhang and Turner, 2008; Waterman et al., 2019). Even though differences between artificial and true herbivory have been largely highlighted in literature (e.g., Baldwin, 1990; Reymond et al., 2000, Lehtilä and Boalt, 2008), mechanical wounding still stands out as the most useful and frequently employed tools to decipher the mechanisms underlying herbivore-induced responses in plants. In fact, numerous ground-breaking discoveries into the molecular aspects of the plant immune system were revealed by subjecting plants to mechanical wounding, including the dynamics of local and systemic responses to insect damage, the action of plant hormones such as jasmonates (JAs) and ethylene in the induction of defence responses and the role of host-cell derived molecules as endogenous immune signals (Green and Ryan, 1972; Creelman et al., 1992; Constabel et al., 1995; O’Donnell et al., 1996; Li et al., 2002; Zhang and Turner, 2008; Fiorucci et al., 2022). These collective findings provide a long and lasting impetus to utilize mechanical wounding as a method to elucidate the consequences of mechanical stimulation in plants.

Tomato (*Solanum lycopersicum*) is the most important horticulture crop in the world and a valuable model to study plant genetics and development (Carvalho et al., 2011; The Tomato Genome Consortium, 2012; Rohan et al., 2019). Tomato has classically served as a remarkable tool to uncover fundamental aspects of the plant defences to mechanical stimulus, including artificial and true herbivory, serving as a practical and useful system to understand the signal transduction events leading from injury to activation of defence responses (Green and Ryan, 1972; Farmer and Ryan, 1990; Constabel et al., 1995; Li et al., 2004; Campos et al., 2009; Wang et al., 2018). However, on an economical perspective, the production of this vegetable is still considered of high risk due to its remarkable susceptibility to diseases and pests, which cause severe losses in fruit quality, nutritional value, and yield (Gibertson and Batuman, 2013; Wang et al., 2021). This scenario indicates that, even though we have a robust knowledge on how tomato anti-herbivore traits are activated in response to biotic attack, a more holistic perspective is necessary to fully grasp the consequences of the mechanical stimulation caused by herbivory on plant development and to eventually develop methods aiming to mitigate the results of such stressful condition in our agroecosystems.

Here we demonstrate how mechanical wounding affects the overall development of tomato plants. Following the classical paradigm of “growth versus defence”, which invokes the existence of a physiological trade-off where activation of growth suppresses defence responses and vice-versa (Herms & Mattson, 1992; Huot et al., 2014: Sestari & Campos, 2022), we observed that repetitive lesions on tomato leaflets impairs the formation of multiple growth traits, including shoot elongation and flower formation, while concomitantly activating the production of defence responses such as trichome formation and the expression of anti-herbivory-related genes. We also found that this wound-induced growth reduction is dependent on jasmonates (JAs), a class of plant hormones broadly studied for its regulation of the growth versus defence antagonism (Yang et al., 2012; Leone et al., 2014; Campos et al.; 2016; Guo et al., 2018a,b; Fiorucci et al., 2022). Besides providing a mechanistic perspective on how tomato plants adjust their development when subjected to a specific environmental stimulus, our results may offer a practical explanation for why tomato productivity is so negatively impacted by herbivores that wound green tissues to meet their nutritional demands.

## Material and methods

### Plant material and growth conditions

For all described experiments, plants were kept in a growth room at 26 °C (± 0.8 °C), 66 % (± 10 %) relative humidity under 16 h at a light intensity of 250 μM m^−2^ s^−1^ and 8 h dark. Tomato (*Solanum lycopersicum*) seeds, cultivars Micro-Tom (cv. MT, kindly donated by prof. Lazaro E. P. Peres – Universidade de São Paulo, Brazil; Carvalho et al., 2011) and Santa Clara (commercially available from Feltrin Sementes ® – Farroupilha, Brazil), were germinated on plastic pots containing Carolina Soil (Carolina Soil ® – Santa Cruz do Sul, Brazil). Fourteen days after germination, seedlings were transplanted to individual pots also containing Carolina Soil. Irrigation was performed daily by supplying water to the trays containing the pots. Selection of *jasmonate insensitivei1-1* (*jai1-1*) homozygous plants was performed following a PCR-based genotyping protocol described by Li et al. (2004), with primers described in Supplemental Table S1. Apart from the experiment shown in the Supplemental Figure S1, where the cv. Santa Clara was used for estimation of growth rate, all data was obtained using the cv. MT.

### Mechanical wounding

Twenty-five-day old tomato plants were subjected to mechanical wounding with the aid of a haemostat with serrated tip, following the protocols described by Zhang and Turner (2008) and Herde et al. (2013), with small adaptations. The haemostat tip was vigorously forced across the tomato leaflets to produce clearly visible lesions that cross the midvein (Figure 1A). All available true leaves were wounded at least once in a sequential manner, from the bottom to the top of the plant, and from the terminal to the basal leaflets, for a total of ten unique wounds per plant. Differently from described in the Zhang and Turner (2008) protocol, all ten lesions were performed in the same day.

**Figure 1.**
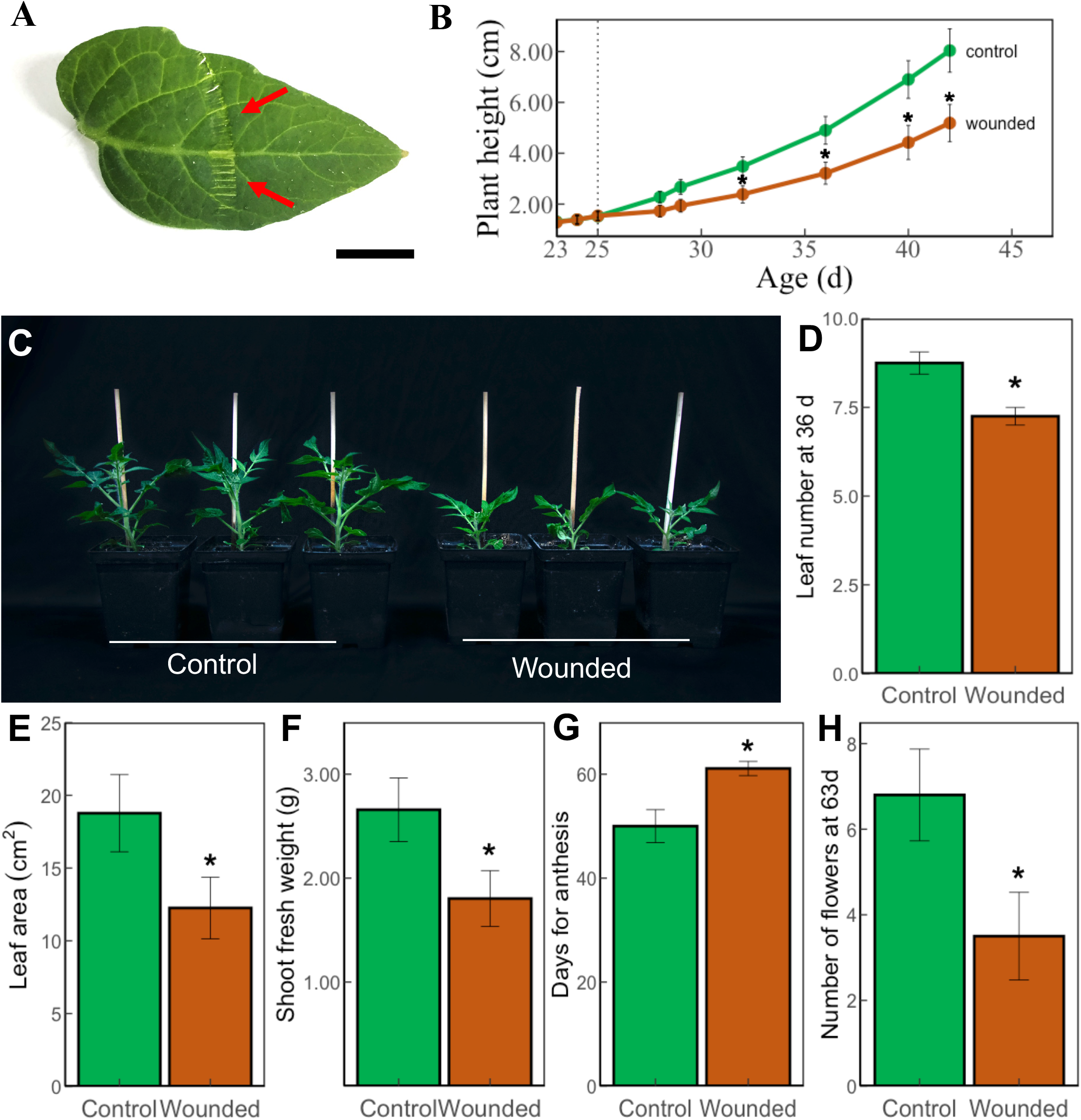
Effect of mechanical wounding on tomato development. (A) Tomato (cv.MT) leaflets were repetitively wounded using a serrated haemostat that was firmly clamped across the tissue to cross the midvein and produce clearly visible lesions, as indicated by the red arrows (bar = 1 cm). (B) Effect of wounding on tomato growth rate. Lesions were performed when plants were 25-d old, as indicated by the dotted line. (C) Representative individuals of 42-d old control and wounded plants (17 DAW). (D) Leaf number of 36-d old (11 DAW) and (E) leaf area of 45-d old (20 DAW) control and wounded plants. (F) Shoot fresh weight of 45-d old (20 DAW) control and wounded plants. (G) Average days for anthesis and (H) number of flowers per plant in 63-d old (38 DAW) control and wounded individuals. Data in all graphs represent the mean ± s.e.m of at least 8 plants. Controls represent unwounded plants. Asterisks denote significant differences between control and wounded plants at p<0.05 (Student’s t-test).

### Evaluation of growth and flowering parameters

Before lesions were performed, plants with similar height and leaf number were pre-selected to reduce variation among the population. Growth rate of control (unwounded) and wounded plants was determined by measuring shoot length with a measuring tape and by counting the number of true leaves formed during time. The same set of plants was subsequently used to assess the number of days taken to open the first flower (days for anthesis) and the number of flowers formed at the 63-d timepoint. Estimation of leaf area was performed using ten fully expanded leaves taken from the middle section of 45-d old plants. Leaves were photographed and the resulting images were used to calculate total area using the ImageJ software (Schneider et al., 2012). Shoot fresh weight was evaluated by immediately weighting the excised aerial parts (without roots) of 45-d old plants in a precision scale. The material was then dried in an oven for five days and weighted again to estimate shoot dry weight.

### Evaluation of defence parameters

Two traits directly related to resistance against herbivorous arthropods were evaluated as an indication of the activation of defence parameters in tomato: the density of leaf glandular and non-glandular trichomes and the expression of the anti-digestive and defence-related genes *Proteinase inhibitor I* (*PI-I*) and *Proteinase inhibitor II* (*PI-II*) (Li et al., 2004; Campos et al., 2009).

Trichome density was evaluated by marking a 0.25 cm^2^ area in the terminal leaflet and counting every trichome found using a dissecting microscope. Tomato trichomes were identified based on stalk length and format and the presence and absence of terminal glands, according to Simmons and Gurr (2005). Since trichome density in tomato is dependent on leaf age (Li et al., 2004), measurements were performed on fully expanded leaves taken from the middle section of 45 d-old plants. At that age, leaves that were previously wounded are all localized in the bottom section of the plants, thus not being utilized for trichome quantification.

For gene expression analysis, RNA was extracted from control and wounded leaflets of 26-d old plants, 24h after wounding. Frozen leaflets were homogenized with a mortar and pestle and total RNA was extracted from using a RNeasy kit (QIAGEN) with on-column DNase (QIAGEN) treatment. cDNA was reverse transcribed using 1μg total RNA with Superscipt First Strand Synthesis System for RT-PCR (Invitrogen) with 18mer oligo-dT. All steps were performed following manufacturers protocols. Quantitative real-time amplification (qPCR) from reverse transcribed samples were conducted in a RotorGene 3000 thermocycler (Corbett Life Science, Australia) using primers for the tomato *PI-I* and *PI-II* genes (Supplemental Table S1). Evaluation of gene expression was performed using the method described by Livak and Schmittgen (2001), normalizing the transcript levels to the tomato *Glyceraldehyde phosphate dehydrogenase* (GAPDH – Zhong & Simons, 1999). Three independent RNA samples (biological replicates) containing a pool of leaflets from three different plants were evaluated per treatment.

### Statistical analysis and data availability

Statistical inferences were made using Student’s *t*-test at the 5% level of significance, always comparing the treatments and genotypes to the MT control (unwounded). All experiments were independently repeated three times with similar results, using a minimum of 10 plants per treatment (unless otherwise noted). All data supporting the findings of this study are available from the corresponding author upon request.

## Results

### Mechanical wounding hinders tomato development

To evaluate the effects of mechanical lesions on tomato development we have adapted simple and reproducible protocols that are commonly utilized to study wound responses in the plant model *Arabidopsis thaliana* (Zhang and Turner, 2008, Herde et al., 2013). Briefly, 25 d old tomato plants (cv. MT) were wounded using a serrated haemostat that was firmly clamped across the leaflets, perpendicular to the midvein, to produce clearly visible lesions (Figure 1A). At this age, MT plants usually have two to four true leaves, all of which were wounded in a consecutive manner (see methods) for a total of ten wounds per plant. Measurements of shoot growth rate indicate that, as observed for *A. thaliana* (Yan et al., 2007; Zhang and Turner, 2008; Fiorucci et al., 2022; Shi et al., 2022), repetitive mechanical wounding significantly reduces the size of tomato plants (Figures 1B-C). Despite statistical significance in growth reduction (p<0.05, according to Student’s t-test) was only observed seven days after wounding (7 DAW, 32 d of age), shoot growth rate changed at early time points (3 and 4 DAW, 28 and 29 d of age, respectively) when compared to unwounded control (Figure 1B). At 42 d of age (17 DAW), shoots of wounded plants were 35,6% shorter than those of control plants (Figure 1B). Similar results were obtained when mechanical wounding was performed in a commercially non-miniature tomato, cv. Santa Clara (Feltrin Sementes - Brazil), where lesions also begin to significantly reduced shoot growth rate 7 DAW (32 d of age) when compared to control unwounded plants (Supplemental Figure S1).

Reduction in shoot stature is not the only developmental process that is negatively affected by mechanical wounding in tomato. We also observed that lesions hinder the capacity of MT plants to develop new leaves, which is indicated by a significant reduction in the number of leaves per plant at 36 d (11 DAW, Figure 1D) and in the average leaf area at 45 d (20 DAW, Figure 1E) in wounded plants. Wounded plants also accumulate about 30% less aerial biomass (g of fresh weight) compared to control unwounded plants at 45 d (Figure 1F). Interestingly, no significant alteration (Student’s t-test, p<0.05) in plant dry weight was observed (Supplemental Figure S2), suggesting that lesions may negatively affect water uptake or water retention in plants. Finally, wounding delayed tomato flowering as wounded plants took approximately 10d longer to open their first flower (Figure 1G) and developed 51% less flowers than control unwounded plants at 63 d (Figure 1H). Taken together, our findings indicate that mechanical wounding hinders numerous growth and flowering traits in tomato plants.

### Wounding-induced growth reduction in tomato is dependent on jasmonate (JA) signalling

Several lines of evidence indicates that plant responses to mechanical stimulation are largely dependent on JAs. For instance, JA levels are enhanced when plants are subjected to mechanical wounding (Creelman et al., 1992; Glauser et al., 2008; Pandey et al., 2017) and mutants impaired in JA signalling fail to activate responses to lesions (Yan et al., 2007; Chung et al., 2008; Zhang and Turner, 2008). To test if the wound-induced growth reduction observed in tomato is dependent on the JA signalling, we compared wounding responses of the tomato *jai1-1* mutant to the MT wild type plants. *jai1-1* carries a genetic lesion in the tomato homolog of CORONATINE INSENSITIVE1, an F-box protein that is an essential component of the JA receptor in plants (Li et al., 2004). For this reason, multiple JA-related responses are defective in *jai1-1* (Li et al., 2004; Campos et al., 2009). While wounded MT plants display an apparent reduction in shoot elongation when compared to unwounded control MT plants, *jai1-1* control, and wounded plants demonstrate strikingly similar growth rates (Figure 2A). This indicates that, as described for *A. thaliana*, the wound-induced growth reduction in tomato is dependent on a functional JA signalling pathway (Yan et al., 2007; Zhang and Turner, 2008; Campos et al., 2009). Interestingly, we noticed a tendency for *jai1-1* plants to produce slight longer shoots than MT, even in the absence of mechanical stimulus. Even though this is a very subtle phenotype, with statistical differences evidenced only at a single time point (36-d – Figure 2A) and no visual differences observed in *jai-1* and MT control plants during the whole experiment (e.g., at 45-d-old plants – Figure 2B), it is tempting to speculate that JAs may act as endogenous negative regulators of growth in tomato. In fact, similar results have been described in the literature for other plants such as *A. thaliana* and rice, where genetic mutations leading to impairment of the JA biosynthesis and signalling pathway led to plants with promoted growth traits such as longer hypocotyls, longer petioles, and early flowering (Yan et al., 2007; Yang et al., 2012; Major et al., 2017).

**Figure 2.**
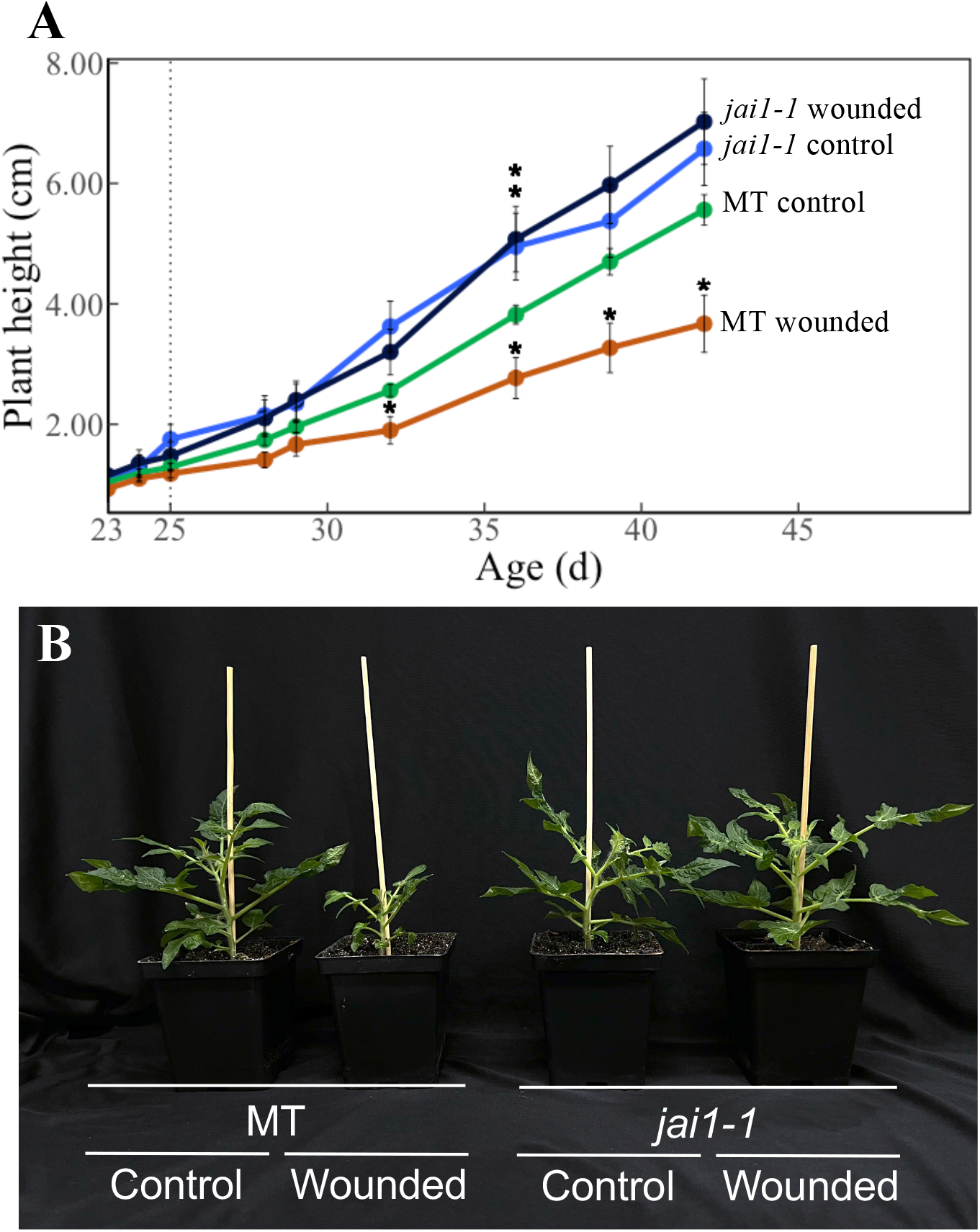
Tomato wound-induced growth repression is dependent on the jasmonate pathway. (A) Effect of wounding on the growth rate of MT (wild-type) and *jai1-1* mutant plants. Lesions were performed when plants were 25-d old (dotted line). Data represent the mean ± s.e.m of at least 8 plants and asterisks denote significant differences when compared to MT control treatment (Student’s t-test, p<0.05). (B) Representative individuals of 45-d old MT and *jai1-1* control and wounded plants (20 DAW). Data in all graphs represent the mean ± s.e.m of at least 8 plants. Controls represent unwounded plants.

### Mechanical wounding promotes the activation of defence-related traits in tomato

The classical growth versus defence paradigm indicates the existence of a growth versus defence trade-off in plants, where growth reduction may be a pleiotropic effect of the promotion of defence responses or vice-versa (Herms & Mattson, 1992; Huot et al., 2014; Ballaré and Austin, 2019; Sestari & Campos, 2022). Thus, to evaluate if the reduced growth induced by our protocol of consecutive mechanical wounding is associated with activation of defence responses in tomato, we quantified the production of two defence-related traits in this species, the density of leaf trichomes and the expression of the defence-related genes *PI-I* and *PI-II* (Li et al., 2004; Campos et al., 2009).

Cultivated tomato (*S. lycopersicum*) possesses five types of trichomes which are usually classified as glandular or non-glandular, based on their length, number of cells and the presence of terminal glands (Simmons and Gurr, 2005). Quantification of trichome density performed on 45-d-old plants (20 DAW), indicates that mechanical wounding promotes the formation of almost all trichomes types in both adaxial and abaxial sizes of newly formed tomato leaves (Supplemental Table S2). When compared to unwounded control plants, wounded plants produce three times more glandular trichomes and ~75% more non-glandular trichomes in their abaxial leaf faces (Figure 3A). A similar result was observed in the adaxial leaf face, where we observed an increase of 103.5% and 63% in the density of glandular and non-glandular trichomes, respectively, in wounded plants when compared to control (Figure 3B). These results are consistent with the described for *A. thaliana*, where artificial wounding also increases trichome density in leaves (Traw and Bergelson, 2003).

**Figure 3.**
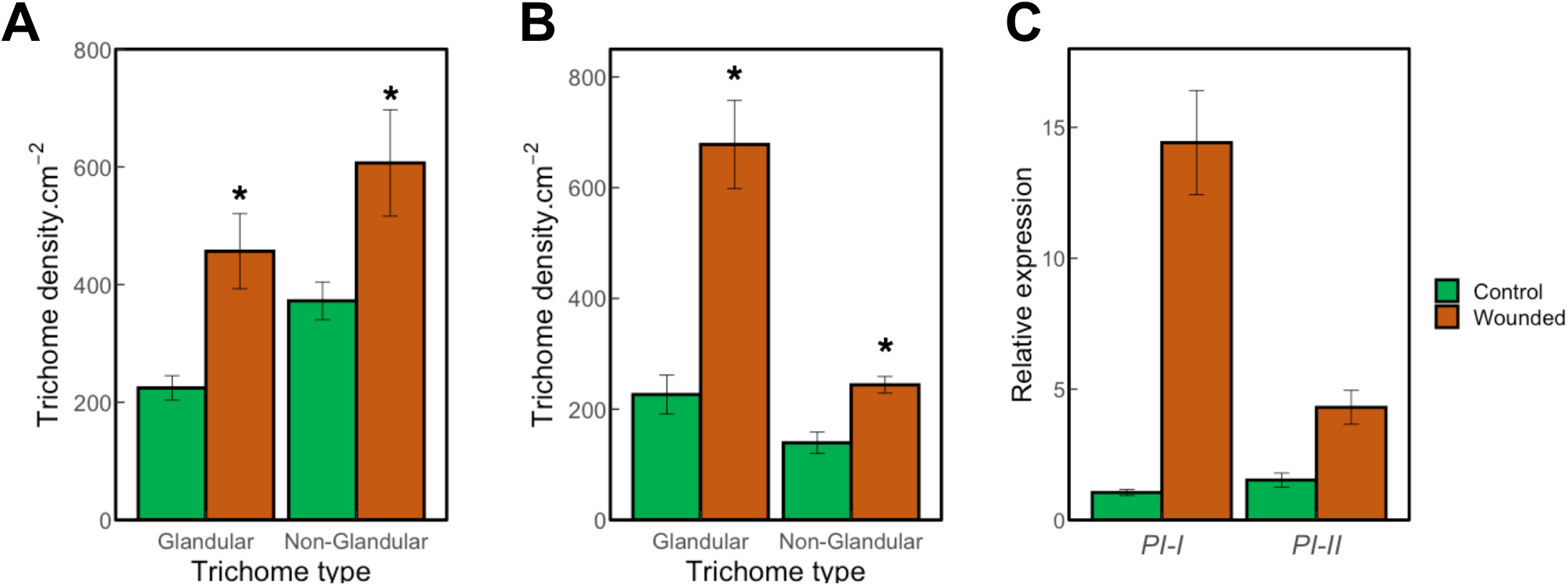
Mechanical wounding induces defence responses in tomato. Effect of mechanical wounding on the formation of glandular trichomes (types I, VI and VII) and non-glandular trichomes (types III and V) on the abaxial (A) and adaxial (B) leaves of 45 d-old tomato (20 DAW). Data represent the mean ± s.e.m of at least 10 plants. Asterisks denote significant differences between control and wounded plants at p<0.05 (Student’s t-test). (C) qPCR analysis of *PI-I* and *PI-II* gene expression in control and wounded tomato leaflets. RNA was extracted using leaves of 26 d-old plants (24 h after wounding). *PI-I* and *PI-II* expression was normalized according to *GAPDH* transcript levels. Controls represent unwounded plants.

Next, we used qPCR to evaluate how wounding affects the expression of anti-herbivory related *PI-I* and *PI-II* genes. Quantification of gene expression performed on 26-d-old plants using RNA extracted from control or wounded leaves (1 DAW) indicates that this type of mechanical stimulation strongly upregulated the transcript levels of both genes (Figure 3C), as we observed an increase of 1300% and 330% in *PI-I* and *PI-II* levels, respectively, in wounded leaves when compared to controls (Figure 3C). Even though the pattern of upregulation in *PI-I* and *PI-II* gene expression and protein production in response to lesions has been thoroughly documented in tomato (e.g., Green and Ryan, 1972; Li et al., 2002; Campos et al., 2009) this is, to our knowledge, the first description of how these genes behave when plants are submitted to repetitive mechanical wounding.

In sum, our results demonstrate that mechanical wounding disturbs the growth versus defence balance in tomato, promoting the activation of anti-herbivore-related traits (trichome density and *PI-I* and *PI-II* gene expression levels) while hindering the formation of development traits (plant growth and flowering parameters).

## Discussion

The plant growth versus defence antagonism is physiological trade-off classically known for its profound impacts in plant development, which is now being recognized as one of the main factors controlling fitness and the genotypic composition of plants and its enemies in natural and agronomical ecosystems (Herms and Mattson, 1992; Huot et al., 2014; Züst et al., 2012; Bally et al., 2015; Fernandez et al., 2021). Efforts to understand the ecological advantages and the molecular framework regulating growth and defence indicate that this antagonism is context-dependent, where plants continuously monitor environmental inputs to adjust the balance between growth and defence as a strategy to optimize fitness for the specific environmental conditions in which they are subjected (Cipollini et al., 2014; Guo et al., 2018a; Ballaré and Austin, 2019; Sestari and Campos, 2022). However, several gaps remain in our understanding of how these environmental signals are integrated by the plant to regulate growth and defence processes concomitantly. For instance, it is not clear how the perception of specific danger inputs regulates the transition from growth- to defence-oriented development and the intensity of the external signal that should be perceived by the plant to reach a “physiological threshold” where the activation of defences may start to impact growth (or vice-versa). Our results shed some light in these topics, showing how a specific type of mechanical stimulus, the consecutive mechanical wounding performed at the early stages of tomato lifespan, induces this species to develop a morphological and physiological state where the plants active defence responses while hindering multiple growth-related traits. We demonstrate that this so-called “defence syndrome” (Ballaré and Austin, 2019) is remarkable for its rapid responses in the plant, such as the increased expression of *PI-I* and *PI-II* genes a few hours after wounding (Figure 3C), and for it long-lasting effects, including the delay of flowering processes observed more than a month after the lesions were inflicted (Figures 1G-H). These observations may provide a feasible theory for why the productivity of tomato (and other important crops) is so negatively impacted by biotic stressors (Gibertson and Batuman, 2013; Savary et al., 2019), as tissues lesions caused by pest and pathogens may cause similar durable and detrimental responses to growth as the ones observed when subjecting plants to artificial wounding.

One important finding of our work was that the wound-induced growth repression in tomato is mediated by a class of signalling molecules, the JAs. The main evidence for this was the observation that, when we subjected a tomato mutant insensitive to JA (*jai1-1*) to mechanical wounding, no differences in shoot growth were observed when compared those to unwounded control plants (Figures 2A-B). These results suggest that the wound-induced growth repression is not a direct consequence of overall physiological disorders that may occur in damaged tissues (e.g., a reduction in photosynthesis parameters caused by crushing part of the leaflets), but rather by the plant sensing the injuries and activating the JA signalling pathway, which in turns, arrest shoot growth. JAs have been described as potent growth inhibitors in numerous plant species, including tomato, and this class of plant hormones is among the main regulators of the wound-induced defence syndrome in *A. thaliana* (Li et al., 2004; Zhang and Turner, 2008; Fiorucci et al., 2022). While previous work indicate that the tomato *jai1-1* is defective in the formation of defence-related traits, including trichome formation and wound-mediated upregulation of *PIs* (Li et al., 2004; Campos et al., 2009), using this mutant we were able to provide direct evidence that endogenous JAs regulate tomato growth in response to external stimulus. Taken together, these results further reinforce the current idea that JAs evolved as central modulators of the growth versus defence antagonism in plants (Yang et al., 2012; Leone et al., 2014; Campos et al., 2014; Campos et al.; 2016; Guo et al., 2018a,b; Fiorucci et al., 2022).

Tomato has long been utilized as a model system to study defence responses in plants and there is an extensive literature on the effects of wounding as a potent inducer of defence responses in this species (reviewed in Wasternack et al., 2006). On the other hand, while wound-induced growth repression has been documented for other plant species (Zhang and Turner, 2008; Engelberth and Elgelberth, 2019) few descriptions of the consequences of mechanical stimulation to tomato overall development have been shown to this date (Stankovic and Davies 1998; Bhatia et al., 2005). We speculate that this discrepancy is caused by the intensity of the wounding stress applied to the plants, as fewer than ten lesions applied at a single timepoint may not be sufficient to produce an obvious disruption of the growth versus defence balance in tomato. Indeed, it has been long demonstrated that an increasing number of wounds trigger a more robust defence response in tomato plants (Green and Ryan, 1972), which may, in turn, generate a more impactful and easily detectable trade-off to development. This hypothesis may also be expanded to other plant species, as the frequently documented wound-induced growth repression in *A. thaliana* is observed under protocols that cause substantial damage to the leaves, in a similar manner to those applied in our experiment (Yan et al., 2007; Zhang and Turner, 2008; Fiorucci et al., 2022; Shi et al., 2022). Interestingly, the observation that gentle types of environmental inputs such as rain, wind, touch, and even sound waves can arrest plant development (Braam and Davies, 1990; Waterman et al., 2019; Matsumara et al., 2022) may indicate that plants utilize different signalling cascades to perceive and differentially respond to specific types of mechanical stimuli.

Our results also demonstrate that the wound-induced growth repression in tomato does not depend on the genetic background of the plants utilized, as similar results were obtained in two different tomato cv. with different physical architecture (a miniature and non-miniature one – Supplemental Figure S1). In fact, preliminary data obtained in our laboratory may indicate that this response is conserved among the Solanaceae family members, as other species belonging to this group (e.g., eggplant, and wild tomato species) also demonstrated reduction in growth-related traits when these plants were subjected to mechanical wounding (data not shown). These observations also support the now well accepted conception that the cv. MT can be employed, with numerous benefits, as a tool to study growth and hormone interactions in tomato (Marti et al., 2006; Campos et al., 2009; Campos et al. 2010; Carvalho et al.; 2011; Kobayashi et al., 2014).

In conclusion, we here show that mechanical wounding stunt tomato growth (in a JA-dependent manner) while concomitantly activating defence responses. For millions of years plant have been co-evolving with an astonishing number of arthropod herbivores capable of inflicting mechanical damage to the green tissues (Grimaldi and Engel, 2005). In this scenario, the wound-induced disturbance in the balance between growth and defence possibly serves as an ecological strategy that evolved to make plants more resilient while less conspicuous to a potential aggressor. As our agronomical system depends on fast plant growth to achieve the desired productivity, understanding how tomato and other crop species adjust this growth versus defence antagonism in response to mechanosensory stimulus is a fundamental step to mitigate the negative consequences of herbivory and to eventually achieve more reliable food systems worldwide.

## Acknowledgements

The authors would like to thank Beatriz C. Araújo for support during *jai1-1* genotyping experiments and Dr. Javier E. Moreno for critical comments on the manuscript.

## Authors contribution

A.F.A.C., P.H.D.R. and M.L.C. designed research, A.F.A.C., P.H.D.R., A.C.A., V.J.P. and D.G.S.P. performed the experiments, A.F.A.C., P.H.D.R. and M.L.C. analysed the data and A.F.A.C. and M.L.C. wrote the manuscript.

## Abbreviations

cv: cultivar
DAW: days after wounding
*GAPDH*: *Glyceraldehyde phosphate dehydrogenase*
JA: jasmonate
*jai1-1*: *jasmonate insensitivei1-1*
MT: Micro-Tom
*PI-I*: *Proteinase inhibitor I*
*PI-II*: *Proteinase inhibitor II*

---

**Supplemental Figure S1.** Mechanical wounding negatively impacts the growth of a commercially available non-miniature tomato. Effect of mechanical wounding on shoot growth of tomato cv. Santa Clara. Lesions were performed on day 25, as indicated by the dotted line. Control represents unwounded plants. Data represent the mean ± s.e.m of at least 10 plants per treatment. Asterisks denote significant differences between control and wounded plants at the same age (p<0.05 according to Student’s t-test).

**Supplemental Figure S2.** Effects of mechanical wounding on tomato shoot dry weight. According to Student’s t-test (p<0.05), no significant alteration was observed when comparing the shoot biomass (mg) of 45-d old (20 DAW) control and wounded plants. Data represent the mean ± s.e.m of at least 10 plants per treatment. Control represents unwounded plants.

**Supplemental Table S1.**
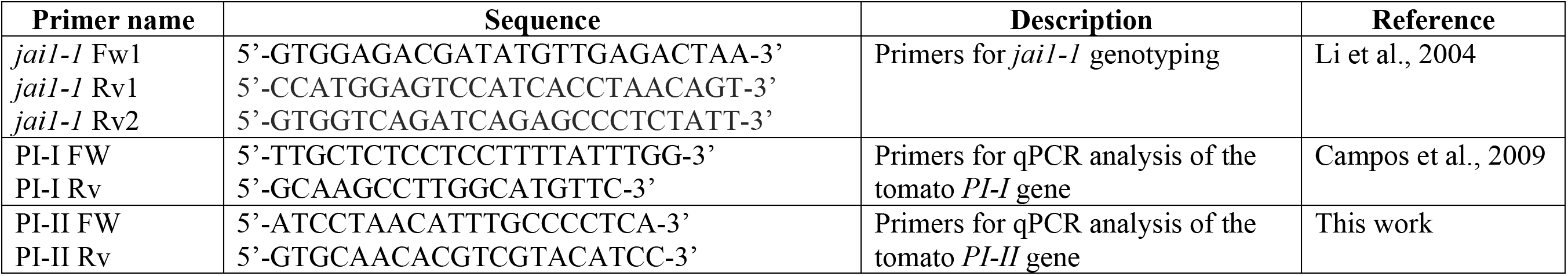
Description of the primers used in the experiments.

**Supplemental Table S2.** Density of tomato trichomes in control and wounded plants. Trichomes were classified according to Simmons and Gurr (2005). Data represent the mean ± s.e.m of at least 10 plants. Asterisks denote significant differences between control and wounded plants for the same leaftlet face at p<0.05 (Student’s t-test). Control represents unwounded plants.

| Treatment | Leaflet face | Trichome type per leaflet face (average.cm-2 ± S.E.M) | | | | |
| --- | --- | --- | --- | --- | --- | --- |
|  |  | I | III | V | VI | VII |
| Control | Adaxial | 5.09 ± 2.59 | 10.18 ± 3.02 | 129.09 ± 18.78 | 205.45 ± 31.99 | 16.00 ± 3.24 |
| Wounded | Adaxial | 8.33 ± 0.44 | 19.33 ± 0.12* | 224.67 ± 12.11* | 620.33 ± 77.31* | 49.33 ± 9.72* |
| Control | Abaxial | 1.82 ± 1.13 | 9.09 ± 2.16 | 363.27 ± 32.16 | 203.27 ± 20.12 | 19.27 ± 2.34 |
| Wounded | Abaxial | 7.33 ± 1.76* | 16.33 ± 3.50 | 590.33 ± 91.13* | 396.33 ± 52.44* | 53.00 ± 12.31* |

